# TreeFix-TP: Phylogenetic Error-Correction for Infectious Disease Transmission Network Inference

**DOI:** 10.1101/813931

**Authors:** Samuel Sledzieski, Chengchen Zhang, Ion Mandoiu, Mukul S Bansal

## Abstract

**Background:** Many existing methods for estimation of infectious disease transmission networks use a phylogeny of the infecting strains as the basis for transmission network inference, and accurate network inference relies on accuracy of this underlying evolutionary history. However, phylogenetic reconstruction can be highly error prone and more sophisticated methods can fail to scale to larger outbreaks, negatively impacting downstream transmission network inference. Additionally, there are no currently available methods which are able to use within-host diversity to improve phylogenetic reconstruction.

**Results:** We introduce a new method, TreeFix-TP, for accurate and scalable reconstruction of transmission phylogenies based on an error-correction framework. Our method uses intra-host strain diversity and host information to balance a parsimonious evaluation of the implied transmission network with statistical hypothesis testing on sequence data likelihood. The reconstructed tree minimizes the number of required disease transmissions while being as well supported by sequence data as the maximum likelihood phylogeny. We use a simulation framework for viral transmission and evolution to demonstrate that TreeFix-TP improves phylogenetic accuracy and downstream transmission network accuracy. We also use real data from ten HCV outbreaks and demonstrate how error-correction improves source detection.

**Conclusions:** Our results show that using TreeFix-TP can lead to significant improvement in transmission phylogeny inference and that its performance is robust to variations in transmission and evolutionary parameters. Our experiments also demonstrate the importance of sampling multiple strain sequences from each infected host for accurate transmission network inference. TreeFix-TP is freely available open-source from https://compbio.engr.uconn.edu/software/treefix-tp/.

## Background

The study of infectious disease has benefited greatly from advances in computational molecular epidemiology. The efficacy of public health efforts to combat the spread of these pathogens has rapidly expanded as technology improves – most notably, the onset of powerful high throughput or next-generation sequencing (NGS) methods has provided molecular epidemiologists with the ability to quickly and cheaply sequence the genomes of the infecting strains (viral or bacterial) [1] which in turn has opened the door for computational analysis of these sequences and of disease transmission. By understanding disease transmission, those investigating a disease can more effectively combat its spread. Computational methods for molecular epidemiology have had a positive impact on public health in a number of cases [2, 3], and continue to be used for the study of infectious disease transmission [4].

Transmission network inference is a challenging computational problem, which has been reflected in the number of new methods developed for understanding disease transmission, especially that of rapidly-evolving RNA viruses [5–10]. A key challenge with studying the transmission of rapidly evolving RNA and retroviruses [11] is that they exist in the host as “clouds” of closely related sequences. These strain variants are referred to as *quasispecies* by virologists [12–16], and the resulting genetic diversity of the strains circulating within a host has important implications for efficiency of virus transmission, virulence, disease progression, drug/vaccine resistance, etc. [17–21]. The advent of next-generation sequencing technologies, has revolutionized the study of quasispecies, but most existing transmission network inference methods are unable to make use of the ability to sequence multiple distinct strain sequences per host. In fact, to the best of our knowledge, there is only one published method, Phyloscanner [7], that explicitly considers multiple strain sequence per host.

Some of the most powerful and widely used techniques for transmission network in-ference, including Phyloscanner [7], are based on computing and using phylogenies of the infecting strains [5–8,22]. We refer to these strain phylogenies as *transmission phylogenies*. These phylogeny-based methods infer transmission networks through a host assignment for each node of the transmission phylogeny, where this phylogeny is either first constructed independently or is co-estimated along with the host assignment. Leaves of the transmission phylogeny are labeled corresponding to the host from which they are sampled, and an ancestral host assignment is then inferred for each node/edge of the phylogeny. This ancestral host assignment defines a transmission network, where transmission is inferred along any edge connecting two nodes labeled with different hosts. In the case of a rooted phylogeny, this coloring also confers direction of transmission, where the host for the ancestral sequence along a transmission edge is considered to be the source of the transmission, and the host of the child sequence is considered to be the recipient. This is illustrated in Figure 1.

**Figure 1.**
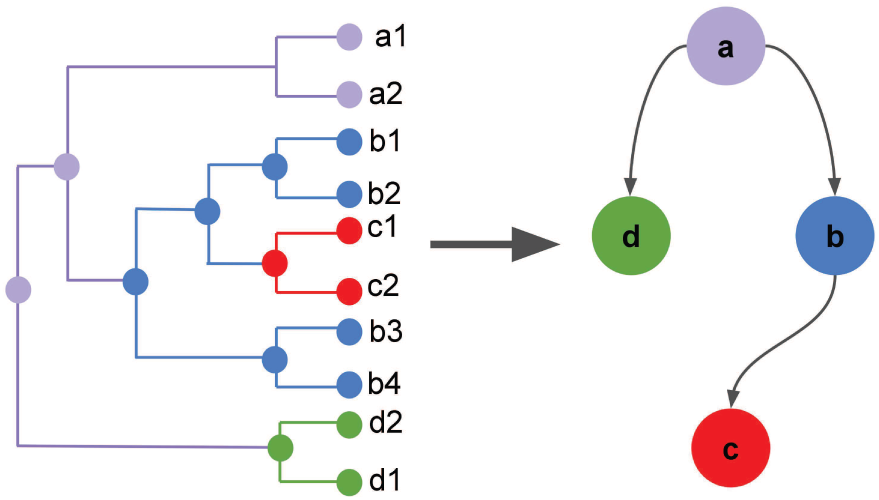
Phylogeny-based transmission network inference: In this figure, internal nodes of the phylogenetic tree on the left are labeled by one of hosts *a, b, c*, or *d*, represented here by the different colors. This labeling of internal nodes causes some of the edges in the tree to have different labels at their two end points, and such edges represent transmission edges in the final transmission network. In the figure we see transitions from *a* to *b, a* to *d*, and *b* to *c*, yielding the transmission network shown on the right.

Two of the most widely-used methods for inference of transmission phylogenies are BEAST [23] and RAxML [24]. For instance, among existing transmission inference methods, TransPhylo [6] uses BEAST to infer a transmission phylogeny, while Phyloscanner [7] uses RAxML. BEAST uses Markov Chain Monte Carlo (MCMC) to estimate phylogenies and evolutionary parameters for several sophisticated models of evolution. Because the models implemented are highly complex, BEAST is prohibitively slow for use on anything other than small data sets. RAxML uses a randomized hill-climbing algorithm to construct a maximum likelihood tree from aligned sequences and is more scalable, but lacks the more sophisticated evolutionary models available in BEAST. As a result it is not considered to be as accurate as BEAST for reconstructing infectious strain evolution. There are also several methods which address transmission phylogeny reconstruction specifically from a transmission perspective, and use transmission information to inform phylogenetic inference. These methods often perform co-estimation of both the transmission phylogeny and network, and often model within-host evolution. BEASTlier [5] and Phybreak [8] both use Bayesian inference for co-estimation of transmission phylogeny and network, and so run into the same scalability issues as BEAST. Thus, even though accurate reconstruction of the transmission phylogeny has a direct impact on transmission network inference, all existing phylogenetic inference methods for transmission phylogenies are either prohibitively slow and unscalable or suffer from poor inference accuracy. Furthermore, none of these existing phylogenetic inference methods can take advantage of the information provided by multiple sequences from each infected host.

In this work, we introduce *TreeFix-TP*, a new method for reconstructing transmission phylogenies that is as scalable as RAxML but significantly more accurate. TreeFix-TP improves the accuracy of infectious disease transmission phylogenies using an error-correction approach. Specifically, TreeFix-TP leverages both sequence and host information to reconstruct more accurate phylogenies than maximum likelihood on its own by minimizing the number of inter-host transmissions while maintaining statistical support. Similar error correction approaches have been successfully used for reconstruction of gene trees [25, 26]; however, these previous methods are based on leveraging a known species phylogeny to error-correct and improve gene trees, and they are therefore inapplicable to the current setting where the goal is to reconstruct the strain tree itself (analogous to the species tree). We address this problem by leveraging intra-host strain diversity and defining a fitness function based on minimizing the number of inter-host transmissions implied by the underlying phylogeny.

In this study, we evaluate the performance of Treefix-TP for transmission phylogeny inference on real and simulated data sets, and compare accuracy of phylogenetic reconstruction to RAxML [24]. We show that TreeFix-TP is robust to variations in transmission model, sequence length, rate of evolution, and number of viruses. Furthermore, we show that using the error-corrected phylogeny as the basis for transmission network inference leads to improved inference accuracy, and demonstrate the use of TreeFix-TP for improving source detection in 10 real-world HCV outbreaks. Finally, we use Phybreak [8] to perform network inference using only a single sequence from each infected host, to evaluate the usefulness of multiple sequences for transmission network inference.

## Methods

### Minimizing inter-host transmissions

The availability of multiple strain sequences from each host provides valuable additional information that can be used to improve the inference of transmission phylogenies. Consider an ideal evolutionary scenario with a complete transmission bottleneck and no reinfection. In such a scenario, all sequences sampled from the same host should form a single monophyletic clade. For *N* hosts, this ideal case would result in a coloring with *N* single-color sub-graphs and would imply *N* − 1 transmissions. Deviations from this ideal would be reflected in the transmission phylogeny and imply a few additional transmissions. Thus, when multiple strain sequences are available from each host, a biologically meaningful criterion for estimating the “correctness” of a transmission phylogeny is to minimize the number of implied inter-host transmissions. Note that the problem of computing the minimum number of implied inter-host transmissions on a given transmission phylogeny is equivalent to the well-known small parsimony problem in phylogenetics and can be solved very efficiently [27]. By minimizing the number of inter-host transmissions implied by a candidate phylogeny, and carefully avoiding over-fitting, we can improve the accuracy of a given phylogeny.

### Description of TreeFix-TP

TreeFix-TP takes as input a multiple sequence alignment of infectious disease sequences, a maximum likelihood phylogeny constructed on the infection disease sequences, and a mapping between all known hosts to the sequences. TreeFix-TP aims to find the transmission phylogeny which is well supported by sequence data and has the minimum transmission cost. Using the maximum likelihood phylogeny as a starting point, we perform iterative local searches and evaluate each candidate tree using a statistical likelihood test and an evaluation of the transmission cost. Candidate phylogenies which are statistically equivalent to the maximum likelihood phylogeny, and with a lower transmission cost, are accepted and set as the starting point for the next local search iteration.

TreeFix-TP uses the Shimodaira-Hasegawa (SH) statistical likelihood test [28] to determine sequence support for a given phylogeny. This test considers two trees, in our case the maximum likelihood phylogeny and a candidate phylogeny, with the null hypothesis that the two trees are equally supported by the sequence data. The null hypothesis is rejected at a significance level *α* which can be defined by the user. If the null hypothesis fails to be rejected, the two trees are considered to be statistically equivalent

The transmission cost for a candidate phylogeny is calculated by solving an instance of the small parsimony problem using Fitch’s algorithm [27]. The states at the leaves of a candidate phylogeny are the hosts from which each sequence is known to be sampled. Fitch’s algorithm, then, calculates the minimum number of state changes required to generate the given phylogeny, which corresponds to minimizing the number of inter-host transmissions. In this case, we are concerned only with the cost of a candidate and not the internal assignments of hosts, so only the upward step of Fitch’s algorithm is performed.

### Algorithmic details

The input for TreeFix-TP is a multiple sequence alignment *A*, maximum likelihood phylogeny *T*_*ML*_, and a mapping *M* : *L*(*T*_*ML*_) → *H*, where *H* is the set of infected hosts and *L*(*T*_*ML*_) denotes the set of leaves of *T*_*ML*_. The output is a transmission phylogeny *T* *. We perform a local search in a manner similar to TreeFix and TreeFix-DTL [25, 26]. Using *T* ′← *T*_*ML*_ as a starting point, we set *C*′ to be the transmission cost of *T* ′, and a candidate tree *T* is proposed by performing random NNI and SPR operations on *T* ′. Using the SH test, we find a p-value for the null hypothesis that *T* and *T* ′are equivalent. Then, we calculate the transmission cost *C*(*T*) of *T*. *C*(*T*) is calculated using Fitch’s algorithm for the small parsimony problem, where the states are the possible hosts. Set intersection and unions are done efficiently by implementing each nodes set of potential hosts as a bit set. This allows us to take advantage of bit-level parallelism and use bitwise AND and OR word operations to efficiently calculate the parsimony cost of a given phylogeny. *T* is accepted if *T* is statistically equivalent to *T* ′(*p* < *α*), and has a lower transmission cost than *T*′ (*C*(*T*) < *C*′). Otherwise, *T* is accepted with some predefined probability. If *T* is accepted, we set *T*′ ← *T, C*′ ← *C*(*T*), and perform another iteration of local search. By default, TreeFix-TP runs for 5000 iterations, which is often sufficient for trees with a few hundred leaves. For larger trees, more iterations may be necessary to fully explore the local space. At the end of the local search, TreeFix-TP outputs the tree *T* * which is statistically equivalent to *T*_*ML*_ and has the lowest transmission cost found during the search. If multiple trees were found with the same minimum transmission cost, TreeFix-TP outputs the tree with the highest likelihood.

The search can be modified in a number of ways. The model under which likelihood is calculated, likelihood test used, and significance level *α* can all be changed. By default, we use the *GT R* + Γ model, and the SH-test implemented in RAxML [24] with a significance level of 0.05. A custom transmission cost module can also be specified. One possible application of this is to use epidemiological data to weight certain transmissions higher, which would cause TreeFix-TP to prefer phylogenies where that transmission does not occur. Finally, the user can specify the number of iterations to be performed.

### Evaluation using simulated data sets

#### Data set generation

To evaluate the performance of TreeFix-TP, we generated a number of simulated data sets across a variety of parameters and developed a testing pipeline to compare TreeFix-TP with RAxML (see Figure 2). Our simulated viral data sets were generated using FAVITES [29], a recently developed framework for simultaneous simulation of transmission networks, phylogenetic trees, and sequences. The specific parameters chosen for each simulated data set are described in detail in the next section and in Table 1, but a brief overview of the simulation pipeline follows. First, a contact network was generated using the Barabasi-Albert model [30], and one host was randomly selected to be infected. Transmission was then simulated under one of two different compartmental models, either Susceptible-Exposed-Infected-Recovered (SEIR) or Susceptible-Infected-Recovered (SIR) [31]. Transmission was simulated for a predefined amount of time, or until all hosts were recovered. Once it had been determined which hosts would be infected, internal evolution of the virus was simulated under a logistic-growth coalescent model. Each internal phylogeny was then connected according to transmission to form one full transmission phylogeny. The branch lengths of this phylogeny were then scaled by different factors to simulate different rates of sequence evolution. Finally, sequences were simulated using the GTR + Γ model starting with a real HCV viral sequence from HCV outbreak data (discussed in more detail in the section on “Source recovery in HCV outbreaks”). The GTR rate matrix and gamma parameter were determined by applying RAxML to estimate parameters and construct a phylogeny for real sequences from an HCV outbreak. For evaluation of reconstruction accuracy, we used sequences from all infected hosts by default, but also considered the effect of sampling only a subset of infected hosts on network inference accuracy (section titled “Inference accuracy with missing hosts”).

**Table 1.**
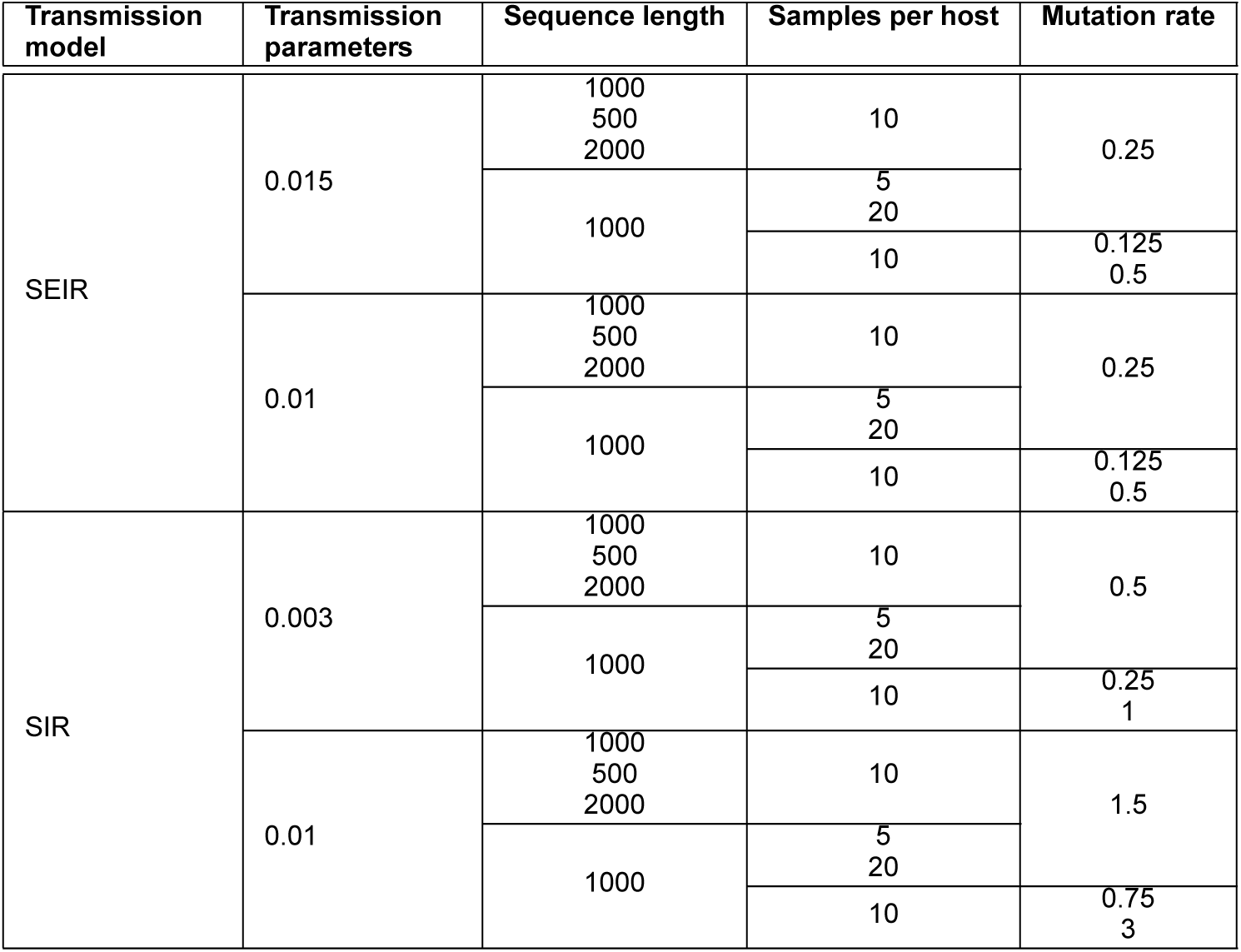
Simulation Parameters: This table shows the set of parameters used for generating the 28 different types of data sets used in our simulation study. For each of these 28 data set types, 20 unique data sets were simulated. These parameters were chosen to mimic realistic viral evolution and to evaluate TreeFix-TP under a wide range of conditions.

**Figure 2.**
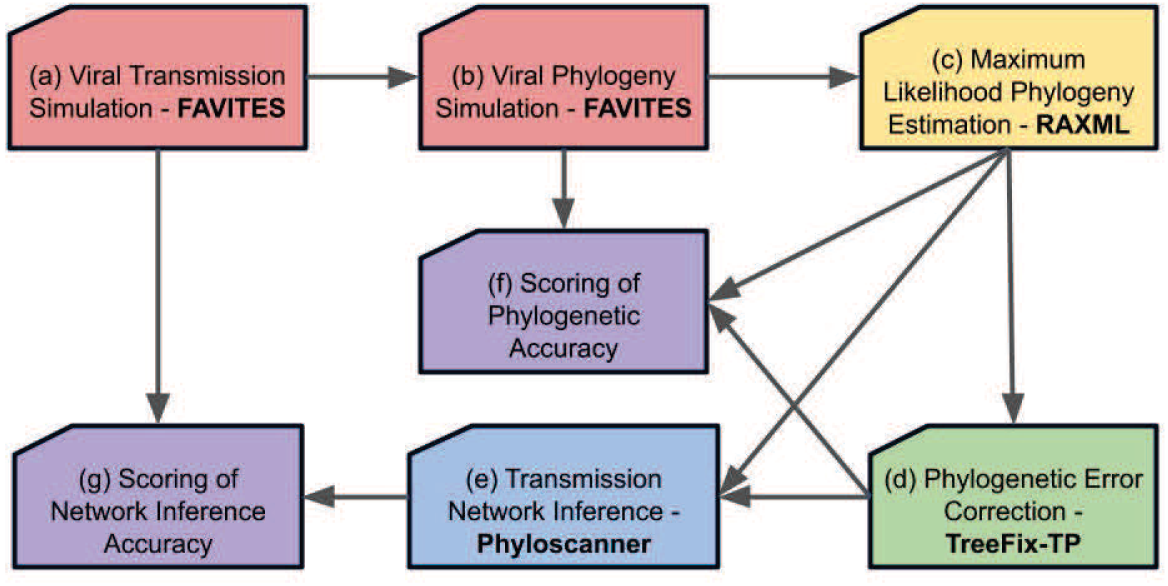
TreeFix-TP Testing Pipeline: To evaluate TreeFix-TP, we first used FAVITES to generate a transmission network (a) and true viral phylogeny (b). Maximum-likelihood phylogenies were then reconstructed from sequences using RAxML (c), and were error corrected with TreeFix-TP (d). The RAxML and TreeFix-TP phylogenies were compared using RF distance, as described in the section on “Evaluating reconstruction accuracy” (f). We then reconstructed transmission networks with Phyloscanner (e), using the RAxML and TreeFix-TP phylogenies, and evaluated the accuracy of the reconstructed networks as described in the section on “Evaluating network inference accuracy” (g).

#### Description of simulated data sets

The simulated contact networks consisted of 1000 individuals, with each individual connected to other individuals through 100 outgoing edges preferentially attached to high-degree nodes under the Barabasi-Albert model. The transmission networks were simulated under both the SEIR and SIR models of transmission. These models are parameterized by transition rates *β, λ*, and *δ*, where *β* is the rate of transition from susceptible to exposed in the SEIR model or susceptible to infected in the SIR model, *λ* is the rate of transition from exposed to infected in the SEIR model, and *δ* is the rate of recovery for infected individuals. In our simulation, we had four categories of data sets with variations on these parameters to explore the effect of transmission model on reconstruction accuracy. Due to the simulation of latent periods, data sets generated under the SEIR model tend to exhibit an outbreak structure, where one high-degree individual infects several of its neighbors, followed by a period of low infection. When one of the newly-infected neighbors becomes infectious, another outbreak occurs. This is contrary to the SIR model, which tends to have a more periodic pattern of disease transmission. In addition to varying the transmission model, we simulated data sets with different rates of infection and recovery. This resulted in four categories of simulation, which we called *SEIR015, SIR003, SEIR01*, and *SIR01* for infection rates of 0.015, 0.003, 0.01, and 0.01 respectively. For most of our results, we group SEIR015 and SEIR01 together, and group SIR003 and SIR01 together. On average, there was no significant difference between the transmission model parameter settings. Parameters for the transmission and within-host coalescent models were chosen so that the within-host phylogeny coalesced close to transmission time, so that there were no clearly long edges separating sequences from different hosts.

By default, we simulated sequences of length 1000 nucleotides and sampled 10 sequences per infected host. We scaled the phylogeny by 0.25 on data sets where the SEIR model was used, and by 1.5 on data sets where the SIR model was used. These scale factors were chosen so that the height of the tree would be approximately ten expected mutations per-site. For each of the four categories, we tested the effects of varying sequence length, number of samples per host, and scale factor, varying one of these parameters at a time from the default setting. Specifically, we simulated sequences of length 250, 500, and 1000, sampled 5, 10, and 20 sequences, and scaled the tree by double or half the default. Including the default setting, this resulted in 7 distinct parameterizations per category, or 28 total. Full details of the parameters for each category can be found in Table 1. Finally, for each set of simulation parameters, we simulated 20 different data sets for a total of 560 simulated data sets. Due to limitations of computation time and availability of memory when running RAxML and TreeFix-TP, we were able to reconstruct phylogenies using TreeFix-TP for 486 of these data sets. Of the 74 runs which did not complete, the simulated trees had an average of 733.43 leaves. The runs were limited to 8GB of memory and 10 days.

#### Basic statistics on data set

For the 486 simulated data sets on which we obtained results, we had between 35 and 630 sequences, with an average of 223.41 leaves. Simulation was restricted to 90 transmissions to limit the tree size, but the number of transmissions was significantly lower for most data sets. The average number of transmissions was 22, and 95% of data sets had between 7 and 49 transmissions. Of the data sets for which we obtained results, only 6 had more than 60 transmissions.

#### Evaluating reconstruction accuracy

The phylogeny reconstructed by TreeFix-TP was evaluated in a number of ways. The accuracy of the reconstructed phylogeny was evaluated by calculating the Robinson-Foulds distance [32] between the true evolutionary history from the simulated data and both the maximum likelihood tree reconstructed by RAxML and the error-corrected tree reconstructed by TreeFix-TP. We calculated the average RF distances, normalized by the maximum possible RF distance (number of internal edges). Additionally, we calculated the *RF percent decrease* as follows: Given simulated tree *S*, maximum likelihood tree *R*, and TreeFix-TP tree *T*, RF percent decrease is given by 100 × (*RF* (*S, R*) − *RF* (*S, T*))*/RF* (*S, R*). Additionally, we looked at the minimum transmission cost implied by the RAxML and TreeFix-TP trees. The cost of the TreeFix-TP tree is guaranteed to be no greater than that of the RAxML tree, but it is valuable to see by how much the transmission cost is decreased and the relationship between transmission cost and Robinson-Foulds distance.

### Evaluating network inference accuracy

Finally, we looked at the accuracy of reconstructed networks as a method of evaluating the impact of TreeFix-TP and improved phylogenetic accuracy on network inference. For each data set, we reconstructed a transmission network with Phyloscanner [7], using either the RAxML and TreeFix-TP trees as input. We chose to use Phyloscanner for network reconstruction because Phyloscanner can infer both a phylogeny and host relationships, but allows us to provide a starting phylogeny. We started with both the RAxML and TreeFix-TP trees, and infer only the relationships between hosts and internal node labeling of the phylogeny. We then interpreted those relationships as a transmission network, and only inferred edges where there was a clear transmission relationship. Phyloscanner labels some internal nodes as unknown, and we did not infer any transmission edges between unknown hosts. We compared these networks to the true simulated transmission network by calculating recall, precision, and F1 score.

We considered reconstructing networks with several other tools such as Phybreak [7], TransPhylo [6], and BEASTlier [5], but only Phyloscanner allowed us to reconstruct our own phylogeny while considering the impact of sampling multiple sequences per infected host. Further, each of these programs had strict constraints on the input data they could handle, so only Phyloscanner was applicable to our data set. TransPhylo models within host evolution, but only considers a single sampled sequence per host. BEASTlier supports multiple sequences, but requires that all sequences from the same host all be clustered together on the phylogeny. Finally, because Phybreak performs simultaneous inference of both phylogeny and transmission network, it was not suitable to evaluate the difference in tree reconstruction methods.

We also evaluated the value of sampling multiple sequences from each infected host by comparing our results to network inference results using only a single sequence. Specifically, we generated networks using Phybreak [8], which co-estimates the phylogeny and transmission network, and compared these networks to the true networks in a similar manner by calculating the average F1 score.

## Results

### Basic experimental results

We obtained results of running RAxML and TreeFix-TP on 486 data sets across a variety of parameters. For our basic test, we evaluated 35 data sets corresponding to the SEIR transmission model, sequence length 1000, 10 sequences per host, and a mutation rate of 0.25. Among these trials, 48.6% of the data sets showed some decrease in RF distance, while 42.86% saw no improvement with the use of TreeFix-TP, and 8.57% saw an increase. The average RF percent decrease for trees which improved was 14.6%, and as high as 46.154%, while the average RF percent increase for those trees that got worse was only 3.644%. In every run where there was no improvement, the maximum likelihood tree generated with RAxML implied exactly as many or only one more transmission than the true number of transmissions, so ability for TreeFix-TP to correct errors by minimizing transmission was limited. Across all 35 data sets, the average normalized RF distance of trees reconstructed with RAxML was 0.152, while trees reconstructed with TreeFix-TP had an average normalized RF distance of 0.137. The overall average RF percent decrease was 6.78%.

We also evaluated 32 data sets corresponding to the SIR transmission model, sequence length 1000, 10 sequences per host, and a mutation rate of either 1.5 or 0.25 (aggregated over both transmission rate categories). The average normalized RF distance of trees constructed with RAxML was 0.103. Trees reconstructed with TreeFix-TP had an average normalized RF distance 0.098. The magnitude of improvement is impacted by the large number of no-change error corrections. Specifically, under the SIR model of transmission, 68.75% of runs had no-change, while 28.13% showed a decrease in RF distance, and the remaining 3.13% showed an increase. The overall average RF percent decrease was 3.66%, but those which improved had an average RF percent decrease of 14.116%, and as high as 28.57%. For those which got worse, the average RF percent increase was 9.8%. A comparison of these results across the SEIR and SIR transmission models suggests that error correction might be more effective under a model of transmission that includes a latent period, which results in transmissions patterns which more closely reflect outbreaks.

#### Impact of varying sequence length

To evaluate the robustness of TreeFix-TP to the amount of sequence information available, we varied sequence length from the base 1000 nucleotides to 250 and 500 nucleotides (Figure 3a). Under the SEIR model, we found that TreeFix-TP continued to improve the accuracy of phylogenetic reconstruction with shorter sequence lengths, and that sequence length didn’t seem to have a large effect on the extent to which error correction improved the accuracy of the phylogeny. At sequence length 1000, the average normalized RF distance decreased by 6.78% from 0.152 to 0.137 after error correction. At length 500, this was a decrease of 10.16% from 0.264 to 0.235. At sequence length 250, the average RF distance decreased by an average of 5.31% from 0.403 to 0.380. Note that, as expected, the absolute error rate increases sharply, for both RAxML and TreeFix-TP, as sequence length decreases.

**Figure 3.**
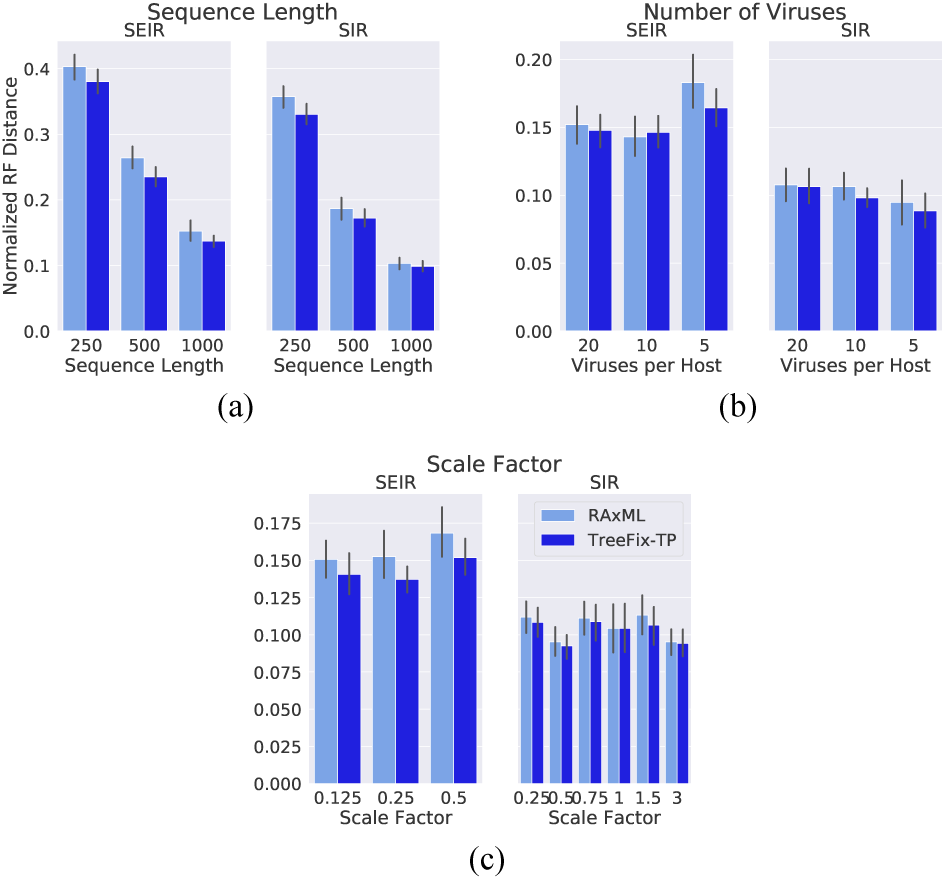
Robustness of phylogeny reconstruction to different parameters: Normalized Robinson-Foulds (RF) distance from the simulated phylogeny for reconstructions with both RAxML and TreeFix-TP under a variety of settings. TreeFix-TP reconstructs the most accurate trees across all data sets. **(a)** RF distance for varied sequence lengths. Trees are in general more accurate with longer sequences, and TreeFix-TP improves upon RAxML to a greater extent with shorter sequences. **(b)** RF distance for varied numbers of viruses sampled from each host. TreeFix-TP has the largest improvement when fewer viruses are sampled. **(c)** RF distance across multiple different scale factors. TreeFix-TP reconstructed the most accurate phylogenies with all scale factors.

Under the SIR model, we found the error correction continued to have an impact at all sequence lengths, and that error correction was more effective at shorter sequence lengths. With sequence length of 1000, the average RF distance decreased by 3.66% (0.103 to 0.099 normalized RF). At length 500, there was a 6.33% decrease (0.187 to 0.172 normalized RF), and at length 250 there was a 7.08% decrease (0.357 to 0.330 normalized RF). Thus, error correction seems to be more effective under this model with shorter sequences, likely because longer sequences contain more information which allows maximum likelihood methods to reconstruct a relatively accurate tree before any error correction occurs.

#### Impact of varying number of viruses

We observed the effect of sampling different numbers of viruses from each infected host, from the default of 10 to 5 and 20 viral sequence samples (Figure 3b). TreeFix-TP reconstructed more accurate phylogenies in each case, with the largest overall improvement occurring for trees with 5 sequences from each host. Under the SEIR model, with 20 viruses, there was an average RF distance decrease of 2.59% (0.152 to 0.148 normalized RF). With 10 and 5 viruses, there were larger decreases of 6.78% and 7.27% respectively (0.152 to 0.137 and 0.183 to 0.164 normalized RF).

Under the SIR model, with 20 viruses, there was an decrease in average RF distance of only 1.56% (0.108 to 0.106 normalized RF). This decrease was 3.66% with 10 viruses and 4.09% with 5 viruses (0.103 to 0.099 and 0.096 to 0.089 normalized RF).

#### Impact of varying scale factor

We found that TreeFix-TP is also robust to various rates of sequence evolution (Figure 3c). Under the SEIR model of evolution, scale factors of 0.125, 0.25, and 0.5 resulted in a decrease in average RF distance by 6.36%, 6.78%, and 7.8% respectively (0.151 to 0.141, 0.152 to 0.137, 0.168 to 0.152 normalized RF). Under the SIR model, we used two different sets of scale factors dependent on the disease transmission parameters. Aggregated across SIR003 and SIR01, we tested scale factors of 0.25, 0.5, 0.75, 1, 1.5, and 3. These scale factors had average RF percent decreases of 2.51%, 2.05%, 2.51%, 0.006%, 5.73%, and 1.003% (0.111 to 0.108, 0.095 to 0.093, 0.111 to 0.109, 0.1043 to 0.1042, 0.113 to 0.106, and 0.095 to 0.094 normalized RF). As expected, the overall RF distances tended to be larger for very small and very large scale factors.

### Network Inference results

To assess the downstream impact of improved phylogenetic accuracy on transmission network inference we used Phyloscanner [7], the only existing phylogeny-based method that can use multiple sequences per host, to reconstruct transmission networks, and then compared them to the true simulated transmission networks. For the 35 baseline data sets under the SEIR model, trees constructed with RAxML led to networks with an average F1 score of 0.645, while networks reconstructed using TreeFix-TP trees had an average F1 score of 0.652. This increase is comprised of increases in recall and precision from 0.578 to 0.584 and 0.751 to 0.759, respectively. We obtained very similar results for the 32 baseline data sets under the SIR model where error correction resulted in an increase in average F1 score from 0.635 to 0.643. This was comprised of increases in recall and precision from 0.559 to 0.567 and 0.758 to 0.763 respectively. Though these improvements in F1 score are relatively small, our experiments clearly demonstrate that improved phylogenetic accuracy does lead to improved transmission network accuracy. However, more sophisticated network inference methods may be needed to fully utilize improvements in phylogenetic accuracy.

As before, we also assessed the impact of various simulation parameters on downstream network inference. These results are described below.

#### Impact of varying sequence length

We evaluated network reconstruction accuracy across sequence lengths of 250, 500, and 1000 nucleotides (Figure 4a). As the figure shows, in most cases, networks generated from TreeFix-TP trees had a higher average F1 score, with improvements in line with what was observed under the baseline data sets (corresponding to 1000 nucleotide sequence length). However, we observed that the improvement was higher for sequence length 250; for instance, under the SEIR model for sequence length 250, there was a 10.69% increase in F1 score from 0.536 using the RAxML phylogeny to 0.600 using the TreeFix-TP phylogeny. This indicates that under certain conditions, especially those where sequence information is lacking, phylogenetic error correction can lead to a significant increase in downstream transmission network inference. As expected, longer sequences tended to result in higher F1 scores overall. The highest average RAxML F1 score with 250 nucleotides was 0.536, compared to 0.663 with 500 nucleotides and 0.652 with 1000 nucleotides.

**Figure 4.**
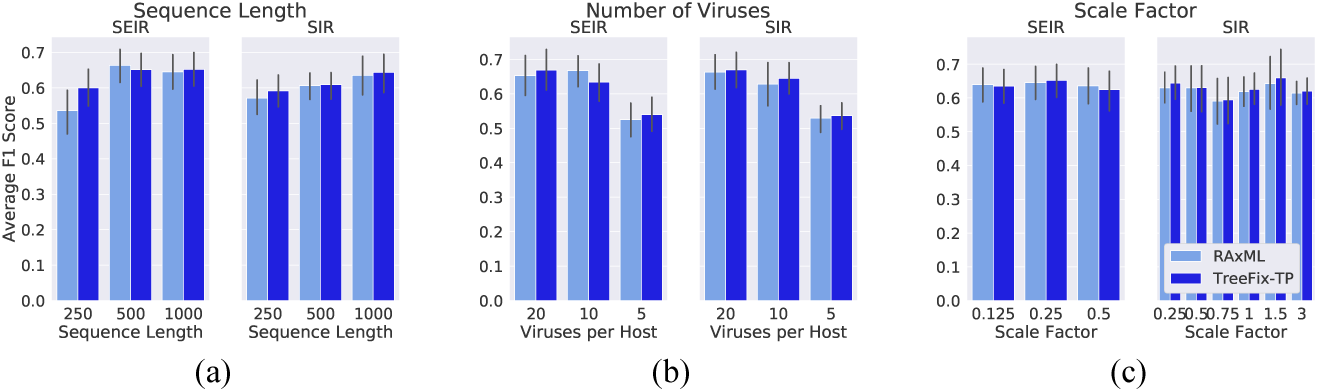
Robustness of Network Inference to different parameters: Average F1 score is shown for networks reconstructed from data sets with varying parameters. TreeFix-TP performs at least as well or better than RAxML under most settings, and is especially useful when less sequence information is available. **(a)** Network reconstruction accuracy for different sequence lengths. Reconstruction tends to be more accurate with longer sequences. **(b)** Network reconstruction accuracy for varying numbers of viral strains sampled from each infected host. Reconstruction tends to be more accurate with more sequences per host. **(c)** Network reconstruction accuracy for different scale factors (mutation rates). There is no clear relationship between scale factor and reconstruction accuracy.

#### Impact of varying number of viruses

Varying the number of viruses sampled from each host, either 5, 10 (baseline), or 20, (Figure 4b) had no significant impact on network inference accuracy. TreeFix-TP resulted in a higher average F1 score for networks in all but 1 case, and the increases were similar to those observed on the baseline datasets (corresponding to 10 sequences per host). The largest improvement seen was a 2.74% increase with 5 viruses per host under the SEIR model. Generally, we found that the overall F1 score increased as more viruses were sampled per host; with 20 and 10 viruses per host, average F1 scores were in the 0.6-0.7 range, compared to 0.5-0.6 with 5 viruses per host.

#### Impact of varying scale factor

As with phylogenetic error correction, scale factor did not seem to have a significant impact on the magnitude of improvement for network inference (Figure 4c). The average F1 scores were highly similar across all scale factors, and there is no clear indication that error correction has a larger impact at any single scale factor. The average F1 score fell between 0.591 and 0.652 for all trials, and the largest different between RAxML and TreeFix-TP F1 scores was 2.67%.

#### Inference accuracy with single sequences

TreeFix-TP takes advantage of sequencing multiple infectious disease strains from each infected host. Likewise, Phyloscanner [7] is designed to work only when multiple strain sequences are available per host. The underlying assumption is that using multiple sequences per host can lead to more accurate inference of transmission networks. To systematically test the validity of this assumption, i.e., to evaluate the usefulness of multiple sequences per host for network inference, we reconstructed networks with a single strain sequence (chosen randomly) per host using the network inference program Phybreak [8] on the 16 baseline SEIR015 runs. Note that Phyloscanner [7] is inapplicable under this single strain sequence setting. We attempted to reconstruct networks using TransPhylo with a dated phylogeny reconstructed by BEAST 2, but were unable to because of excessive runtime. We also attempted to use BEASTlier [5], but it is currently in development for use with BEAST 2 and was prohibitively slow when used with BEAST 1. Phybreak performs simultaneous inference of the phylogeny and transmission network and we ran it for the recommended 50,000 MCMC iterations. We found that using single sequences greatly decreases the accuracy of network inference (Figure 5). Specifically, networks reconstructed using Phybreak had an average F1 score of only 0.16, compared to the 0.57 and 0.59 when multiple sequences were used with PhyloScanner on RAxML trees and TreeFix-TP trees, respectively.

**Figure 5.**
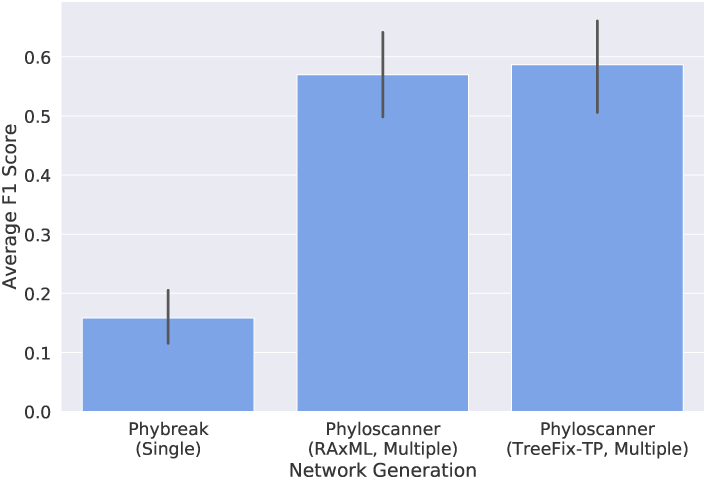
Effect of Multiple Sequences on Transmission Network Inference: Average F1 score of network inference using both a single sequence per host and multiple sequences per host is shown. Networks reconstructed using Phyloscanner with multiple sequences had a significantly higher F1 score, regardless of whether RAxML or TreeFix-TP was used to reconstruct the phylogeny. When only a single sequence per host was used, network reconstruction was much less accurate.

#### Inference accuracy with missing hosts

To simulate the use of TreeFix-TP in disease outbreaks where not all infected hosts have been sampled, we randomly sub-sampled 25, 50, and 75 percent of our hosts, then restricted the sequence data to only those hosts and ran RAxML and TreeFix-TP. We then reconstructed networks using Phyloscanner and evaluated the reconstructed networks in the same way (Figure 6). As expected, the overall average F1 score drops off significantly when data is available from fewer hosts. These experiments also suggest that the improvement in network inference gained by using TreeFix-TP does not carry over to lower levels of host sampling; with 25%, 50%, and 75% levels of sub-sampling, we saw no improvement in network inference when using the error corrected phylogeny. This strongly suggests that TreeFix-TP as well as network inference methods such as Phyloscanner are most effective when there is near-complete sampling of infected hosts.

**Figure 6.**
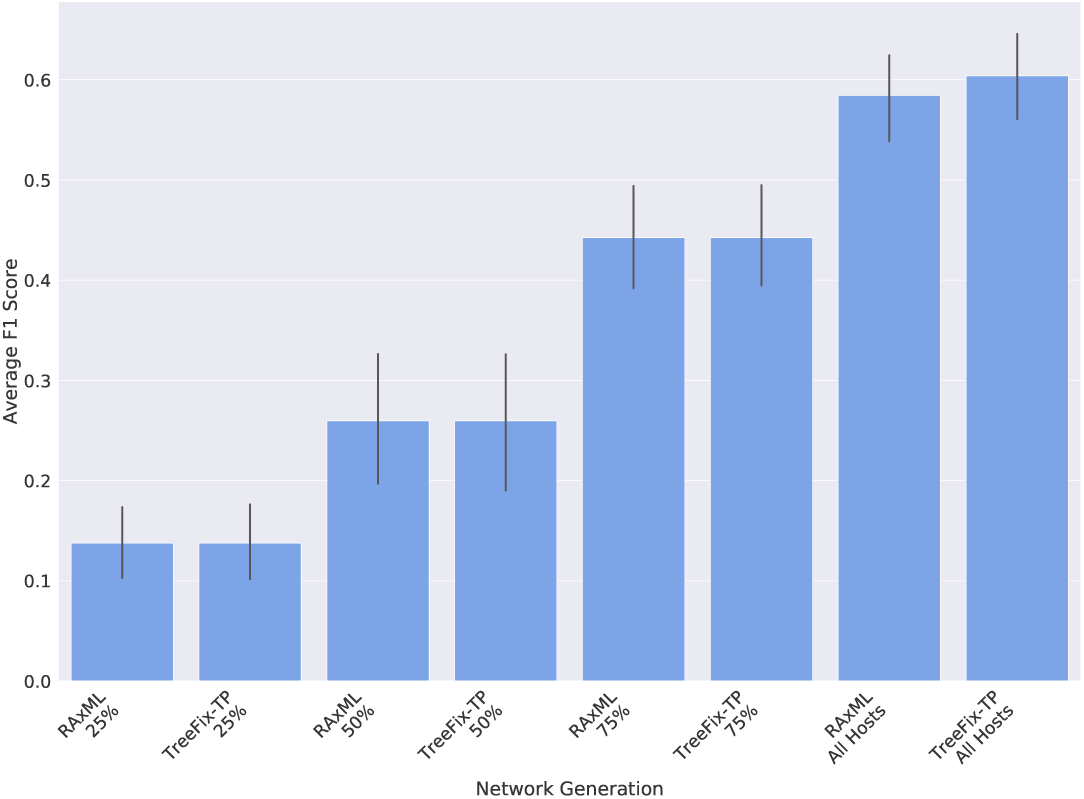
Effect of Missing Hosts on Transmission Network Inference: Average F1 score of network inference is shown when a subsample of hosts is used to reconstruct the phylogeny and networks. Network inference accuracy significantly drops off when not all hosts are sampled from an outbreak. Networks were most accurate when TreeFix-TP was used with all hosts sampled, but the percentage of hosts sampled has a larger effect on F1 score than which tool was used for phylogeny reconstruction.

### Source recovery in HCV outbreaks

We also evaluated the impact of using TreeFix-TP on real data sets of HCV outbreaks made available by the CDC [9]. In total, there are 10 different data sets, each representing a separate HCV outbreak. Each of these outbreak data sets contains between 2 and 19 infected hosts and a few dozen to a few hundred strain sequences. For each of these 10 outbreaks, the source host of the outbreak is known (through the CDC’s epidemiological efforts). We used a simple phylogenetic pipeline to infer a source for each of these 10 data sets as follows: We first constructed phylogenetic trees using RAxML and TreeFix-TP and rooted them using two of the most widely used rooting methods, balanced rooting (implemented in RAxML [24] and midpoint rooting [33, 34]. We then used Sankoff’s algorithm for the small parsimony problem [35] to label the internal nodes of these phylogenies with hosts and reported the host assignment at the root as the inferred source of that outbreak. (Note that Phyloscanner also uses Sankoff’s algorithm to label internal nodes of the phylogeny, but we chose not to use Phyloscanner directly because Phyloscanner is very conservative in its host assignments and often leaves nodes unlabeled.) Using the RAxML trees, the source was correctly identified for 6 and 7 of the outbreaks using balanced and midpoint rooting strategies, respectively. In contrast, the trees reconstructed by TreeFix-TP correctly identified the source in 8 out of the 10 outbreaks with both rooting strategies.

### Running time and scalability

Using its default number of iterations (5000) TreeFix-TP required an average of approximately 37 hours for each run, but this running time varied widely depending on the number of tips and length of sequence. TreeFix-TP took less than an hour and a half for trees of 50-60 tips, but upwards of 200 hours for trees with more than 500 tips and 1000 nucleotide-length sequences. On average, runs took fewer than 9 minutes per tip, and scaled linearly in tree size, number of hosts, and sequence length.

## Discussion and Conclusions

In this paper, we have introduced a new method, TreeFix-TP, for more accurate and scalable reconstruction of infectious disease transmission phylogenies when multiple strain sequences are sampled from each infected host, and demonstrated its impact on phylogenetic inference and downstream transmission network inference. TreeFix-TP uses an error-correction approach where it seeks to improve a given maximum-likelihood phylogeny of the infecting strains by using additional information about which host each strain was sampled from and balancing it with sequence-only likelihood using a statistical hypothesis testing framework. As our experimental results show, TreeFix-TP consistently reconstructs more accurate phylogenies than the state-of-the-art maximum-likelihood phylogeny inference method RAxML. We also showed how TreeFix-TP can be used to augment existing phylogeny-based pipelines for transmission network inference by error correcting the phylogenies before they are used for network inference. As we showed, in most cases the improved phylogenetic accuracy leads to an improved transmission network accuracy. However, there was often only a small improvement in accuracy before and after error correction. This points to the need for more advanced methods for network inference which can take full advantage of improved phylogenetic accuracy.

Going forward, it would be worthwhile to develop even more advanced, yet scalable, methods for construction of transmission phylogenies. As our experimental results show, even though the absolute error rate of TreeFix-TP phylogenies is often significantly lower than that of RAxML trees, this absolute error rate still remains quite high overall even after error correction. This is partly because the ability of TreeFix-TP to error-correct depends on the number of different hosts represented in the phylogeny, rather than on the size of the tree itself. In the future, it may be possible to use additional information about within-host strain evolution to further improve transmission phylogeny inference.

## Ethics approval and consent to participate

Not applicable

## Consent for publication

Not applicable

## Availability of data and material

The simulated data sets used and/or analysed during the current study are available from the corresponding author on reasonable request.

## Authors’ contributions

SS contributed to the theoretical results, implemented the software, performed the experimental study, analyzed the results, and contributed to the writing of the manuscript. CZ contributed to initial project development and conducted the experimental analysis on real data. IM helped supervise the research and contributed to writing the manuscript. MSB conceived the research project, supervised the research, and contributed to the writing of the manuscript.

## Acknowledgements

The authors wish to thank Dr. Pavel Skums (Georgia State University) and the Centers for Disease Control for sharing their HCV outbreak data.

